# Evolutionary constraints in host shifts: limited adaptation of *Plutella xylostella* to cardenolide-defended *Erysimum cheiranthoides*

**DOI:** 10.1101/2025.07.04.663117

**Authors:** Broti Biswas, Erik van Bergen, Teresa Vaello, Laura J. Dällenbach, Christopher W. Wheat, Tobias Züst

## Abstract

1. Plants and herbivorous insects are engaged in long-term coevolutionary arms races, where gains in novel plant defences and corresponding adaptations in herbivores may drive evolutionary change. One such recent innovation is the gain of cardenolide toxins in the Brassicaceae genus *Erysimum*, which resulted in the effective deterrence of most herbivores by these plants. Nonetheless, some herbivores continue to attack *Erysimum*, likely by tolerating cardenolide defences through general detoxification mechanisms that may serve as evolutionary stepping-stones for more specialized resistance.
2. We investigated the interaction between diamondback moth (DBM, *Plutella xylostella*) and its occasional host plant *E. cheiranthoides*, by first screening the standing variation in DBM performance, and second by experimentally evolving DBM populations for increased performance on *Erysimum*. Despite considerable variation among wild DBM populations, larvae consistently avoided *Erysimum* leaves for feeding, and when constrained to *Erysimum*, they exhibited reduced growth rate, survival, and lower adult size compared to individuals feeding on control broccoli plants.
3. Surprisingly, cardenolides could only partly explain the reduced performance of DBM on *Erysimum*, and experimental evolution failed to improve overall performance. Instead, phenotypes of evolved lines converged on what appears to be a pre-existing, highly plastic phenotype found among wild DBM ancestors and which is characterized by rapid development and high weight gain on control plants, but slow development and low weight gain on *Erysimum*.
4. Although DBM failed to evolve improved performance on *Erysimum*, our results demonstrate that its existing genetic variation and phenotypic plasticity are evidently sufficient to support long-term development on *Erysimum*, thereby fulfilling a key condition for the future evolution of specialized adaptations.

## 1. Introduction

Many plants and herbivorous insects are engaged in an ongoing coevolutionary arms race (Ehrlich & Raven, 1964; Marquis et al., 2016). In response to selective pressures from herbivores, many plants have evolved complex defence mechanisms to protect or reduce damage to their tissues. In turn, some herbivores have evolved specialized strategies and counter-adaptations to cope with the defences of their host plant. Repeated cycles of these adaptations and counter-adaptations between plants and herbivores are considered major drivers of genetic and species diversity (Agrawal & Zhang, 2021; Brockhurst & Koskella, 2013).

Key innovations are traits that enable populations to exploit new ecological niches, such as by accessing resources that were previously inaccessible or by improving their ability to avoid natural enemies. Relief from biotic pressures in a novel niche can free populations from ecological and evolutionary constraints, enabling them to expand and diversify rapidly (Miller et al., 2023; Stroud & Losos, 2016). For example, the evolution of extra-floral nectaries (EFNs) in plants facilitated mutualistic interactions with animal defenders, leading to the prediction that higher speciation rates in EFN-carrying lineages compared to sister lineages without EFNs may be due to this key innovation (Weber & Agrawal, 2014). Similarly, host–plant shifts and adaptations linked to detoxification of novel plant compounds have been associated with bursts of diversification in swallowtail butterflies (Allio et al., 2021). Continuous co-evolutionary interactions between plants and herbivores may even lead to repeated and reciprocal key innovations over evolutionary time. For example, plants in the order Brassicales evolved potent glucosinolate defences approximately 85 million years ago (Edger et al., 2015), while Pierid butterflies only began to exploit these glucosinolate-defended Brassicales approximately 10-20 million years later, after they evolved specialized glucosinolate detoxification (Edger et al., 2015; Wheat et al., 2007). Both Brassicales plants and Pierid butterflies have since undergone further evolutionary changes in glucosinolate synthesis and detoxification mechanisms, respectively, with key innovations in each group promoting bursts of speciation (Edger et al., 2015).

Coevolved plant-herbivore systems thus provide compelling evidence for coevolutionary arms races as key drivers of diversification. However, research focusing on coevolved systems generally only allows for retrospective analyses of processes that occurred millions of years ago, and relies on the assumption that present ecological interactions mirror past evolutionary interactions. Furthermore, extant herbivores that co-evolved with their host plants rely on specialized mechanisms to detoxify plant defences, such as the detoxification of glucosinolates by Pierid butterflies (Wheat et al., 2007), or target-site insensitivity, which evolved in monarch butterflies and other milkweed herbivores to tolerate cardenolide toxins (Dobler et al., 2012). Such specialized adaptations likely could only evolve if ancestral herbivore lineages were frequently exposed to the toxic host and had at least a minimal initial tolerance towards this defence. Hence, an improved understanding of plant-herbivore interactions prior to the evolution of key detoxification mechanisms in herbivores is crucial for reconstructing and predicting evolutionary trajectories.

In a recent gain of a defensive key innovation, plants in the Brassicaceae genus *Erysimum* evolved the ability to produce toxic cardenolides, which are co-expressed with ancestral glucosinolates by most species of this genus (Züst et al., 2020). Cardenolides are steroidal compounds that inhibit the activity of animal Na+/K+ ATPase, an enzyme involved in neurotransmission and nerve function (Agrawal et al., 2012). Due to their distinct function from glucosinolates (Blažević et al., 2020), herbivores that specialized on Brassicales by evolving mechanisms to cope with glucosinolate defences are unlikely to be pre-adapted to mitigate the detrimental effects of cardenolides. Indeed, target-site insensitivity, a counter-adaptation that evolved in several herbivores of cardenolide-producing plants, appears to be absent among *Erysimum* herbivores (i.e., Petschenka et al., 2017). Despite the apparent lack of specialized herbivores, *Erysimum* plants are still occasionally attacked by a variety of herbivore species (Mertens et al., 2021), including aphids, bugs, and the diamondback moth *Plutella xylostella* (DBM). The generalized tolerance mechanisms that likely enable these herbivores to cope with cardenolides may therefore serve as an important stepping-stone towards more specialized adaptations.

DBM is a prime candidate for evolving specialized resistance to cardenolides, as it is known to rapidly evolve resistance to the insecticides that are commonly used to protect Brassica crops in agricultural fields (Sarfraz et al., 2005; Sarfraz & Keddie, 2005). DBM feeds on many wild and cultivated plants within the Brassicaceae and is a highly migratory global pest of Brassicaceae crops (Talekar & Shelton, 1993). The species copes with glucosinolates through a specialized adaptation, allowing it to deactivate these compounds by expressing a glucosinolate sulfatase in the fore- and midgut of its larvae, which cleaves the sulphate group from glucosinolates and prevents their activation by myrosinases (Ratzka et al., 2002). In contrast to other *Erysimum* herbivores, DBM populations can easily be maintained in the laboratory, and their short generation times make them well-suited for eco-evolutionary studies, including experimental evolution.

Here we report on a series of experiments to provide novel insights into the coevolutionary interplay between DBM and *Erysimum cheiranthoides* plants. First, we quantify phenotypic and standing genetic variation in twenty DBM populations from distinct geographical locations in Switzerland to *i)* evaluate the fitness costs associated with feeding on *Erysimum* plants, and *ii)* assess whether cardenolide tolerance varies among populations. We then use an experimental evolution approach to *iii)* test if consistent selection pressure to survive on *Erysimum* can drive the evolution of increased DBM tolerance to *Erysimum* and its novel defence in real time.

## 2. Materials and methods

### 2.1. Plant material and cultivation

All population maintenance and experiments were performed using young plants of either *Erysimum cheiranthoides* (*‘Erysimum’* hereafter), broccoli (*Brassica oleracea* var. *italica*), or a fast-cycling variety of *Brassica rapa* (Wisconsin Fast Plants, Caroline Biological Supply Co., USA). For *Erysimum*, we used a pool of seeds from approximately 20 plants collected in a natural population in Northern Germany (53.0669° N, 11.4684° E; ancestors of inbred accession ‘Elbtalaue’, Arabidopsis Biological Resource Center accession CS29250, National Plant Germplasm System accession PI 691911). Large batches of seeds of a broccoli cultivar were purchased from a commercial supplier (‘Coastal Selection Z’, Zollinger Bio, Switzerland), while for fast-cycling *B. rapa*, we used a pool of seeds from a mixed set of accessions maintained at the Department of Systematic and Evolutionary Botany, University of Zürich.

All three species were planted weekly following a standardized schedule for the duration of all experiments in this study, using plastic pots (7 × 7 × 8 cm) filled with peat-based planting substrate (Einheitserde, Patzer Erden GmbH, Germany). Seeds of *Erysimum* were soaked in water prior to sowing and stored at 4 °C for three days to break dormancy. Each pot was sown with three seeds and placed in a growth chamber (constant light, 23°C, 60% RH). After one week, seedlings were removed or transplanted to leave two plants per pot. On day 17 after sowing, *Erysimum* plants were moved to a climate-controlled greenhouse (artificial light supplementation to 16:8 L:D, maintained at 21-23°C, 60% RH), and plants were used for experiments 4-5 weeks after sowing. Broccoli seeds were germinated in moist soil under identical growth chamber conditions. After ten days, seedlings were transplanted in pairs into new pots containing the same planting substrate. On day 17 after sowing, pots were moved to the greenhouse, and plants were used 3-4 weeks after sowing. Young plants of fast-cycling *B. rapa* were used as a neutral oviposition substrate for adult DBMs and initial food for first-instar larvae during experimental setups. Seeds were sown on moist soil and seedlings were transplanted singly into new pots 7 days after sowing. On day 11 after sowing, plants were moved to the greenhouse, and plants were used for experiments 11-18 days after sowing. All plants were checked twice daily and watered as needed.

### 2.2. Establishment and maintenance of 20 DBM populations

Twenty laboratory populations of DBM were established from insects collected at twenty locations across Switzerland between June and August 2021 (Table S1). At each location, late-instar larvae, pupae, and/or adults were collected from *Brassica* vegetable crops. One population (ZH.BOGA) was established from larvae and adults found on *Erysimum cheiranthoides* growing in the Botanical Garden of the University of Zürich. When DBM were collected as larvae in the field, we reared them to adulthood on commercial Chinese cabbage leaves (*Brassica rapa* var. *pekinensis*).

All adults were kept in fabric mesh cages (‘adult cages’ henceforth, 24.5 x 24.5 x 24.5 cm, BugDorm, MegaView Science Co., Taiwan) and provided with a source of 10% sugar water with a cotton wool wick. Adult cages were kept at ambient temperature (19-25°C) under natural light, with supplemental fluorescent lighting on a 16:8 L:D cycle. Small *B. rapa* plants were placed in each adult cage, and after two days of oviposition, egg-covered plants were removed and used to establish the first generations of laboratory-reared DBM.

For the first three generations, egg-covered plants for each population were placed in separate plexiglass cages (20 x 35 x 50 cm) and kept in the same room as adult cages. In this initial phase, newly hatched larvae were fed exclusively on *B. rapa* plants, supplemented with commercial Chinese cabbage when necessary to provide *ad libitum* feeding. From the fourth generation onwards, the larval rearing was switched to broccoli plants as main food source, and large fabric-mesh cages (‘larval cages’ henceforth, 24.5 x 24.5 x 63 cm, BugDorm, MegaView Science Co., Taiwan) were used instead of plexiglass cages for improved ventilation. Larval cages were set up in four identical climate-controlled growth chambers with vertical LED light strips (FitoClima 600 PLH, Aralab, Portugal) set to 23°C, 60% RH, and a 16:8 L:D cycle. Each growth chamber housed a cohort of five populations, staggered in developmental stage by one week, with an initial generation time of five weeks for each cohort.

To initiate a new discrete DBM generation (‘set-up’ hereafter), egg-covered *B. rapa* leaves of each population were added to separate larval cages with three pots (each containing two plants) of 3-week-old broccoli each. Cages were checked and plants watered three times a week, and additional plants were added as necessary. As larval cages only fit nine pots, older (eaten) plants were cut at the base and pushed to the side, so that empty pots could be removed to make space for new plants. Old plant material was left in the larval cage until all DBM had completed development. On average, each population required 30 pots (60 plants) of broccoli to complete larval development.

Adult eclosion began approximately 18 days after set-up. Starting at first eclosion, we removed and counted all adult moths from larval cages three times a week over two weeks, aspirating adult moths with an air pump. Adult moths of each population were added to adult cages, combining repeated collection timepoints of the same population. To avoid overcrowding, we added a maximum of 350 adults to one adult cage over two weeks, discarding excess adults at each collection timepoint when necessary. Given the large spread in timing of eclosion and limited life span adult moths, we collected eggs over four time intervals of 3-4 days each, adding a fresh *B. rapa* plant to adult cages at the start of each time interval. At the end of each interval, plants were removed, and egg-covered leaves were stored for 0-11 days in a fridge at 4 °C before set-up of the next generation. Under these conditions, cold storage inhibits DBM egg development without affecting subsequent development at normal temperatures (Liu et al., 2002). For set-up of the next generation, we used a subset of eggs from all four collection intervals to include a maximum of genetic diversity in the next generation. After final collection of adults, larval cages were frozen for >24h, cleaned, washed with detergent, and soaked in disinfectant (Halamid-D®, Centrachem AG, Switzerland) to limit disease transmission.

### 2.3. Quantifying the costs of *Erysimum* feeding in 20 DBM populations

Larvae from the 20 DBM populations were used to characterize herbivore behaviour and to quantify costs of feeding on *Erysimum* using three types of assays: 1) association (choice) assays which measured the innate preference of naïve first-instar larvae; 2) growth rate assays to quantify short-term weight gain of second-instar larvae on different host plants; and 3), life-history assays to track development of larva from second instar through eclosion, thus allowing us to quantify survival, development time, and adult mass at eclosion on different host plants. Each assay was performed in four blocks staggered by one week each, tracking the set-up of cohorts of the main DBM rearing. Surplus eggs from the peak collection interval of the main DBM rearing were used to prepare experimental larvae for each assay.

#### 2.3.1 Association assays

We used association assays to measure the innate preference of first-instar DBM larvae between leaf disks of 4-week-old *Erysimum* and 3-week-old broccoli plants. Assays were performed on each DBM population 7-11 generations after their initial collection (due to differences in collection time) and repeated once more for each population 1-4 generations later to determine consistency of choice results. We prepared choice arenas in 90 mm diameter Petri dishes with a thin layer of 3% agar, and six leaf disks (9.2 mm diameter) — three from each plant species — were arranged in a hexagonal pattern. Newly oviposited eggs were placed onto a fresh *B. rapa* leaf, and later, 20-25 freshly hatched larvae were transferred to the centre of each choice arena using fine brushes. Larvae were allowed to explore the choice arena for 24 hours, after which we counted the number of larvae feeding on each leaf disc using a stereomicroscope. In both rounds of assays, we set up 7-8 replicate choice arenas per population, resulting in a total of 14 independent choice assays per population across two repeats.

#### 2.3.2 Growth rate assays

We performed growth-rate assays for each population, 6-8 generations after their initial collection. This assay allowed us to accurately track weight gain of individual second-instar larvae over 48 hours, and to include a cardenolide supplementation treatment to isolate and evaluate the specific role of cardenolides, independent of other plant traits. Newly oviposited eggs were added to *B. rapa* plants in custom-made sleeves (8 x 16 x 50 cm) made from fine mesh fabric and kept in the same growth chambers as the main DBM rearing (23°C, 60% RH, 16:8 L:D). After 9 days of development, larvae of comparable sizes were selected from each sleeve and individually weighed using a microbalance (0.01 mg accuracy; Ohaus Explorer EX125, Ohaus Europe GmbH, Switzerland). We selected larvae weighing between 0.4 and 0.7 mg for this assay to maximise comparability of results.

Growth-rate assays were conducted using freshly cut leaf discs (14 mm diameter) placed in 12-well plates (25 mm diameter wells, Greiner Bio-One, Switzerland), with each well containing a thin layer of 1% agar to prevent desiccation of the leaf discs. Each weighed larva was randomly assigned to one of three treatments: a) feeding on a broccoli leaf disc, b) feeding on broccoli with a cardenolide supplement, or c) feeding on a *Erysimum* leaf disc. For the cardenolide supplement, we dissolved the cardenolide digitoxin (Sigma-Aldrich, Switzerland) in 100% methanol at a concentration of 665 µg/mL. We applied 20 µL of this solution to one side of each broccoli disc to achieve a concentration of approximately 3.6 µg of cardenolide per mg of broccoli dry weight. This concentration corresponds to the average cardenolide levels reported for *Erysimum* leaves by Mirzaei et al. (2020). To account for the potential effects of the organic solvent, we added 20 µL of pure methanol to all leaf discs in the other treatments. At application, solvent was spread evenly across each leaf discs and then left to evaporate for 1 hour.

After placing a pre-weighed larva into each well, plates were sealed with parafilm and placed in the growth chamber. Larvae were weighed again after 24 hours and transferred to a second set of 12-well plates containing fresh, identically treated leaf discs. Weights were recorded a third time after an additional 24 hours.

#### 2.3.3 Life-history assays

We conducted life-history assays for each population, 8-11 generations after their initial collection. Following an established protocol (van Bergen et al., 2017), this assay assessed how survival, development time, and adult dry mass were affected by feeding on *Erysimum* plants relative to broccoli plants. As for growth rate assays, newly oviposited eggs of each population were added to *B. rapa* plants in fine-mesh fabric sleeves. After 9 days, 80 larvae weighing between 0.4 – 0.7 mg were selected in each population and randomly distributed among four new sleeves containing two pots (four plants) of 4-week-old *Erysimum* or 3-week-old broccoli plants (20 larvae per sleeve, 2 sleeves per plant species). The sleeves were returned to the growth chamber and watered regularly as needed. Given the low number of larvae added per sleeve, damage was limited, and the initial plants survived throughout the assay. Larvae began to pupate in sleeves after 7 days of development, at which point we began to carefully remove pupae from the sleeves and placed them individually in 1.5 mL Eppendorf tubes with perforated lids for ventilation. The tubes were placed back into the growth chamber and checked daily for adult eclosion. On the day of eclosion, tubes were frozen at -20 °C, and the sex of each adult was later determined by examining their abdominal shape and external genital structures (Figure S1) under a stereomicroscope. Survival was determined at the level of each sleeve as the number of eclosed adults out of the 20 initial larvae. Sexed moths were dried at 60 °C for 48 hours and weighed (0.01 mg accuracy) to determine adult dry mass, and individual development time was determined as days between setup and eclosion.

### 2.4. Experimental evolution of DBM on *Erysimum*

We used an experimental evolution approach to test whether increased performance on *Erysimum* could evolve from standing genetic variation. To maximize genetic diversity at the initiation of experimental evolution, we established a panmictic population by outcrossing all 20 wild DBM populations, 4-6 generations after their initial collection. In brief, the five DBM populations within each cohort were initially subjected to a controlled hybridization scheme, wherein 30 virgin males from one population were crossed with 30 virgin females from a different population, effectively ensuring each population was crossed with two others (Table S2). In the second outcrossing generation, 200 adults from each cross, totalling 1000 individuals per cohort, were pooled into two adult cages and allowed to mate at random. Their offspring — now carrying genetic material from five populations — were again reared to adulthood. In the final outcrossing generation, 1000 adults per cohort (4000 total) were combined in three large ‘larval’ cages and allowed to mate randomly. The resulting eggs formed a fully panmictic ancestral population, from which replicate selection lines were established.

We used batches of approximately 800 eggs from the ancestral population to establish 16 replicate selection lines, eight lines evolving on *Erysimum* and eight lines evolving on broccoli (as cardenolide-free ‘controls’). For experimental evolution we followed the same cohort structure as in main DBM rearing, with each cohort comprising four lines (two Erysimum, two broccoli) staggered by one week. Maintenance of selection lines was mostly identical as for the general DBM rearing, with *Erysimum* selection lines being fed with 4-week-old *Erysimum* plants and requiring approximately 60 plants to complete development. We monitored adult emergence each generation, maintaining a population size of 400–600 individuals to prevent overcrowding while minimizing genetic drift. During the experiment, two populations experienced severe bottlenecks (<100 adults emerging) and were terminated at mid-experiment. To maintain balanced replicate line numbers, two additional populations, together representing a complete cohort, were also removed at this time. Development time of all selection lines accelerated due to adaptation to laboratory conditions and required us to shorten the experimental timing from five to four weeks per generation after 14 generations of selection. Selection was maintained for a total of 25 generations, after which all lines were terminated.

To continuously assess selection line performance, we measured the body mass of emerging adults from all lines each generation. On the day of peak emergence, approximately 30 adults per line were collected and stored at –20 °C. From this sample, individuals were sexed under a stereomicroscope until 10 males and 10 females were selected for each line. Sexed individuals were placed individually into 1.5 ml Eppendorf tubes with pierced lids. Tubes were dried at 60°C for 48 hours and adult mass was determined to the nearest 0.01 mg.

### 2.5. Tracking performance of evolving DBM lines

To assess changes in performance of evolving DBM lines under standardized conditions we conducted the same assays as those used for quantifying performance of wild DBM populations. Due to logistical constraints, not all assays could be performed with the same frequency; however, each assay was performed at least once at both the beginning and the end of experimental evolution. Larval feeding preference was assessed in association assays in generation 3 and generation 26 (using eggs from the final generation under selection). Each choice assay was replicated six times per selection line and generation. Larval growth on broccoli leaf discs, broccoli leaf discs with digitoxin, and *Erysimum* leaf discs was assessed for each selection line using growth rate assays in generations 3, 10, and 26. Finally, larval survival, development time, and adult mass at eclosion on *Erysimum vs.* broccoli were assessed for all selection lines using life-history assays in generations 4, 12, 18, and 24. Two selection lines could not be assessed in generation 24 due to insufficient egg numbers, thus life-history assays were repeated for these lines and two additional selection lines in generation 25. Data for these two generations was pooled for analysis.

### 2.6. Statistical analyses

All statistical analyses were conducted in R, v4.4.3 (R Core Team, 2021), using a combination of linear or mixed effects models, as appropriate. For models with multiple explanatory or fixed variables, we included a complete set of interactions, followed by stepwise removal of non-significant interaction terms. Significance of fixed effect in models with interactions was assessed using Type III Wald chisquare tests implemented in the *Anova* function from the car package (Fox & Weisberg, 2019).

For all association assays, we summarized results of each choice arena as the proportion of larvae choosing *Erysimum vs.* broccoli leaf discs. For the 20 wild DBM populations, we analysed this binary response using a generalized linear model (function *glm*) with a quasibinomial distribution to account for overdispersion. Population, experimental repeat, and an interaction term were fitted as explanatory variables, and significance was assessed using chisquare tests on residual deviance. We then performed a linear regression between population means across the two repeats to estimate an approximate broad-sense heritability (H^2^) in choice behaviour. For experimental evolution lines, we analysed responses across generations using a generalized linear mixed model (function *glmmTMB* in the glmmTMB package) with a beta-binomial distribution (Brooks et al., 2017) to account for overdispersion. Selection line was treated as random effect, while selection regime, generation of selection, and an interaction term were fit as fixed effects.

For growth rate assays, we first estimated relative growth rates (RGR) for each individual larvae using

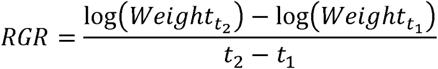

As weights had been measured at three time points, we calculated separate RGR values for the first (0-24h) and second (24-48h) interval. RGR is expected to be constant if larvae were to grow at a constant, near-exponential rate, but exponential growth is often unrealistic for lepidopteran larvae (Tammaru & Esperk, 2007), and larvae additionally experience growth deceleration at the end of each instar (Kivelä et al., 2020). Many larvae in our experiment underwent moult during growth assays, and in fact, RGR values for the first and second interval were negatively correlated (Pearson’s *r* = -0.179, p < 0.001), suggesting that most larvae experienced moult-induced growth deceleration in one of the two intervals. To avoid averaging a non-linear process with few datapoints, we instead used the maximal RGR value estimated for each larva (i.e., RGR of the first or second growth interval). For the 20 wild DBM populations, we analysed RGR_max_ using a linear mixed effects model (function *lmer* in the lme4 package) with assay plate treated as random effect, while treatment, population, and an interaction term were included as fixed effects. We also fit an equivalent model, but treating population as random effect, to estimate broad-sense heritability in growth rates from variance components. For experimental evolution lines, we analysed RGR_max_ using a similar model, including assay plate nested within selection line as random effects, and treatment, selection regime, assayed generation, and all significant interactions as fixed effects.

For life history assays of the 20 wild DBM populations, we first estimated survival at the level of replicate sleeves, using the number of successfully eclosed adults (out of initially 20 larvae) *vs.* the number of dead individuals as a binary response. We analysed survival using a generalized linear model with a binomial distribution, fitting population, assay plant species, and an interaction term as explanatory variables. Development time (time from sleeve setup to eclosion) and log-transformed adult mass at eclosion were analysed at the level of individual moths, using linear mixed effects models (function *lmer* in the lme4 package) with replicate sleeves treated as random effect, and population, assay plant, sex and all significant interactions included as fixed effects. We also fit separate models for all three life-history traits on broccoli and *Erysimum* separately, treating population as random effect to estimate broad-sense heritability in growth rates from variance components.

For experimental evolution lines, we analysed life history data using an equivalent set of models. Survival across generations was analysed using a generalized linear mixed model (function *glmmTMB*) with a beta-binomial distribution to account for overdispersion. Selection line was treated as random effect, while assay plant, selection regime, generation of selection, and all significant interaction terms were included as fixed effects. Development time and log-transformed adult mass were analysed using linear mixed effects models, with replicate sleeves nested within selection line treated as random effects, while assay plant, selection regime, generation of selection, and all significant interaction terms were included as fixed effects.

For wild DBM populations, we compared the estimated population means for each trait and treatment against each other using a set of pairwise correlations. For more straightforward comparison of traits, we used the inverse of development time (i.e., development speed) for higher trait values to reflect better larval performance. Likewise, in association assays we had estimated larval choice as percent larvae feeding on *Erysimum*, but as no larvae exhibited a preference for *Erysimum*, we instead used inverse numbers to reflect strength of *Erysimum* avoidance (and thereby strength of preference for Broccoli) for correlation analyses.

Finally, we had monitored adult performance in each generation of experimental evolution using the weights of a small set of randomly selected, newly emerged adults (10 individuals per sex and generation). We modelled the log-transformed adult dry mass across generations using a linear mixed-effects model (function *lmer* in the lme4 package), including selection line as random effect, and treating sex, selection regime, generation, and all significant interactions as fixed effects. Generation was first treated as factorial variable, but we later repeated the model treating generation as continuous variable when it became clear that temporal trends were approximately linear. Log-slopes of the two selection regimes were compared using the *emtrends* function (emmeans package), while mass at the beginning and end of selection was compare using pairwise linear contrasts (function *emmeans* in the emmeans package).

## 3. Results

### 3.1. Preference and performance of wild DBM populations

All wild DBM populations were assayed once (twice for association assays), 6-11 generations after initial collection. In two-choice association assays, first-instar DBM larvae consistently avoided feeding on *Erysimum* leaves and preferred to feed on broccoli leaves (Figure 1). On average, larvae were twice as likely to be found feeding on broccoli (likelihood of feeding on *Erysimum*: 0.329 ± 0.008, mean ± 1SE). However, population means significantly differed in relative preference (quasi-binomial GLM, χ² = 60.84, df = 19, p = 0.036), with one population (SG. SEVE) showing no significant avoidance of *Erysimum* plants (Figure 1). Relative preferences of populations remained consistent between repeat assays (separated by 1-4 generations, depending on population), indicated by a lack of interaction between population and assay repeat (χ² = 22.84, df = 19, p = 0.910), and resulting in a positive correlation among population means between repeats (approximate broad-sense heritability H^2^: 0.44 ± 0.20).

**Figure 1.**
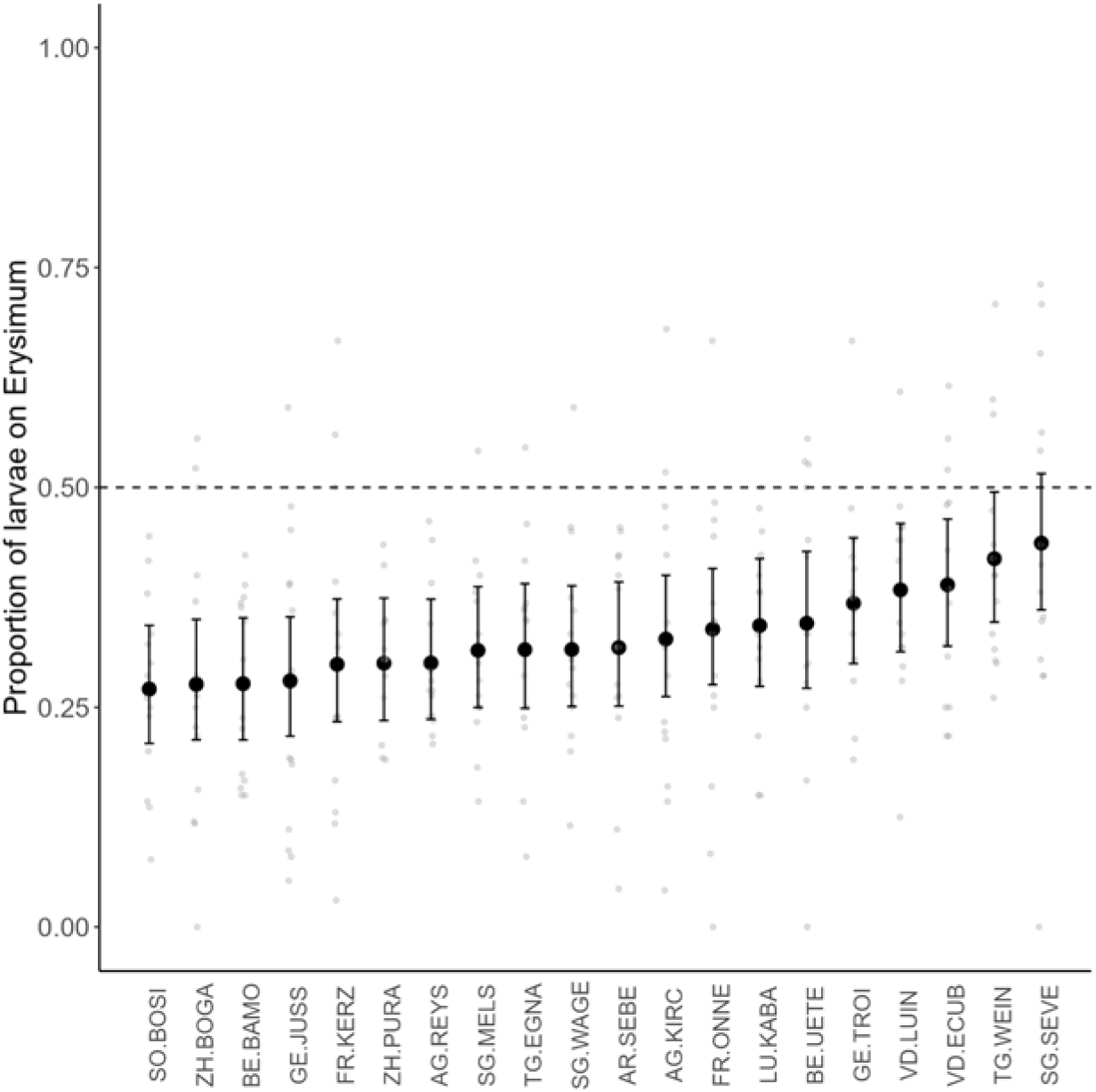
Choice of first-instar DBM larvae feeding on *Erysimum vs*. broccoli leaves in two-way choice assays across 20 wild populations. Populations are ordered by their relative preference for *Erysimum*. Large black dots and error bars are the mean likelihood for each population with 95% confidence intervals, while small grey dots are replicate choice assays (n = 280) of approximately 24 larvae each. The dashed line indicates equal preference for each plant; values <0.5 correspond to avoidance of *Erysimum*, while values >0.5 suggest preference for *Erysimum*. Overlapping of confidence intervals with the 0.5 line indicates a lack of preference at the p = 0.05 significance level.

We next performed growth-rate assays to quantify growth of second-instar caterpillars over a 48-hour period on leaf discs of broccoli, broccoli with added digitoxin, and *Erysimum*. RGR_max_ significantly differed across treatments (χ² = 78.00, df = 2, p < 0.001), with larvae growing fastest when feeding on broccoli leaf discs. Addition of the cardenolide digitoxin reduced maximal growth rate by 8%, while feeding on *Erysimum* leaf discs reduced growth by 17% (Figure 2). RGR_max_ also differed substantially among populations (χ² = 60.09, df = 19, p < 0.001), with larval growth on broccoli ranging from 0.030 ± 0.002 mg mg^-1^ h^-1^ to 0.041 0.002 mg mg^-1^ h^-1^ (mean ± 1 SE; equivalent to 37% reduction from highest to lowest). There was no statistical support for a population-specific response to treatments (χ² = 43.71, df = 38, p = 0.242), suggesting that treatment-imposed reductions affected all populations equally. Despite a significant main effect of population, we only found weak heritabilities for RGR_max_ in the three assay treatments (broccoli: H^2^ = 0.09; broccoli+digitoxin: H^2^ = 0.07; *Erysimum*: H^2^ = 0.06).

**Figure 2.**
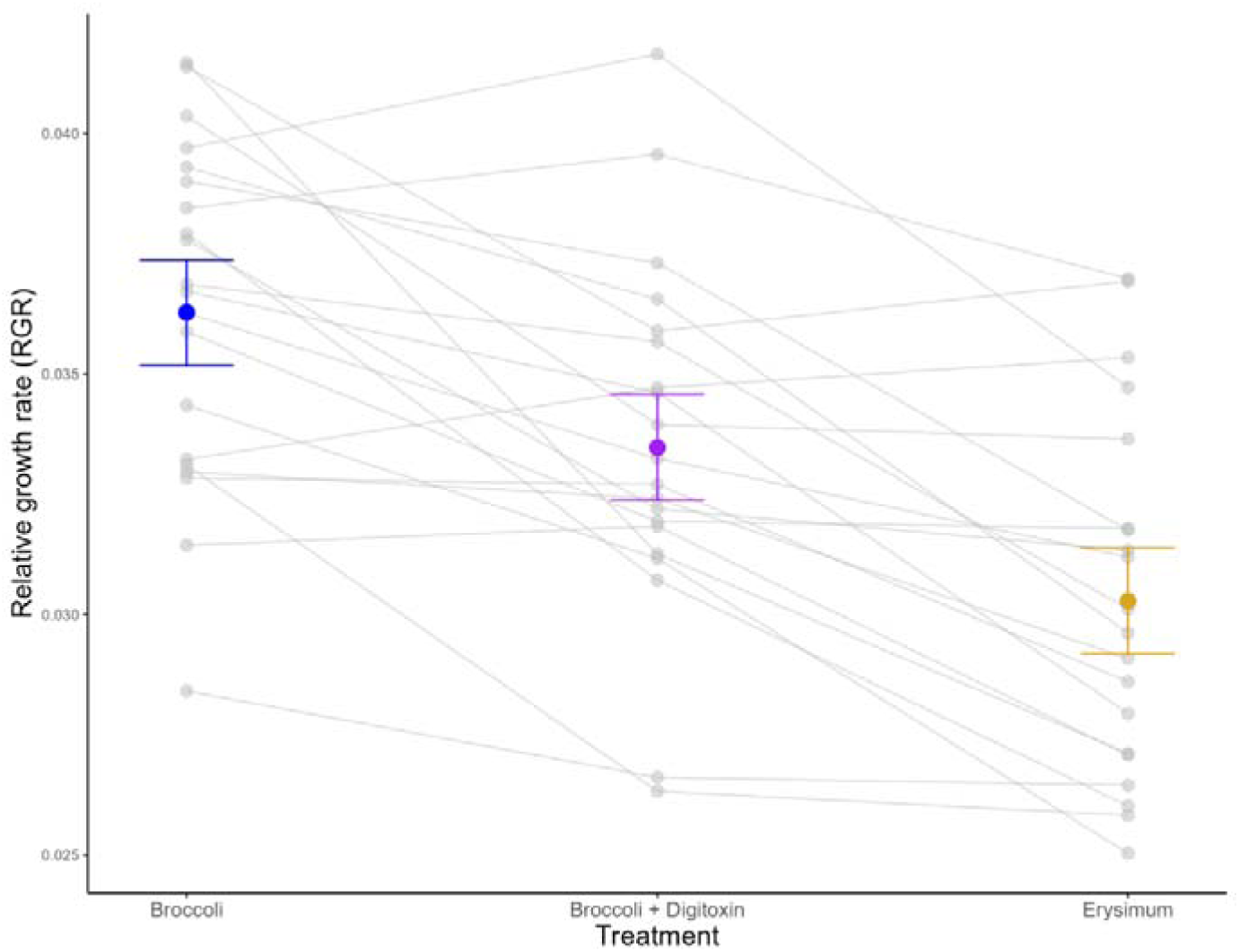
Growth of larvae from 20 wild DBM populations feeding on leaf discs of broccoli (blue), broccoli with added digitoxin (purple), or *Erysimum* (orange) for 48 hours. Growth was estimated as relative growth rate (RGR) over two 24h intervals (0-24h and 24-48h), retaining the higher value for each larva. Coloured dots and error bars are global means of RGR with 95% confidence intervals. Grey points are the mean maximum growth rates of each population across treatments, while grey lines connect means of the same population in each treatment. Note that there was no statistical support for a treatment × population interaction, thus population-specific patterns have to be interpreted with care.

Life-history assays revealed that survival of second-instar larvae to adulthood was 21% lower on *Erysimum* compared to broccoli plants (Figure 3a, χ² = 87.69, df = 1, p <0.001). Survival varied significantly among populations (χ² = 138.65, df = 19, p <0.001), with a weak but significant interaction between assay plant and population (χ² = 33.36, df = 19, p = 0.022). While this suggests population-specific differences survival on the two host plants, heritabilities for mean survival of populations were again low on each host plant (broccoli: H^2^ = 0.09; *Erysimum*: H^2^ = 0.05).

**Figure 3.**
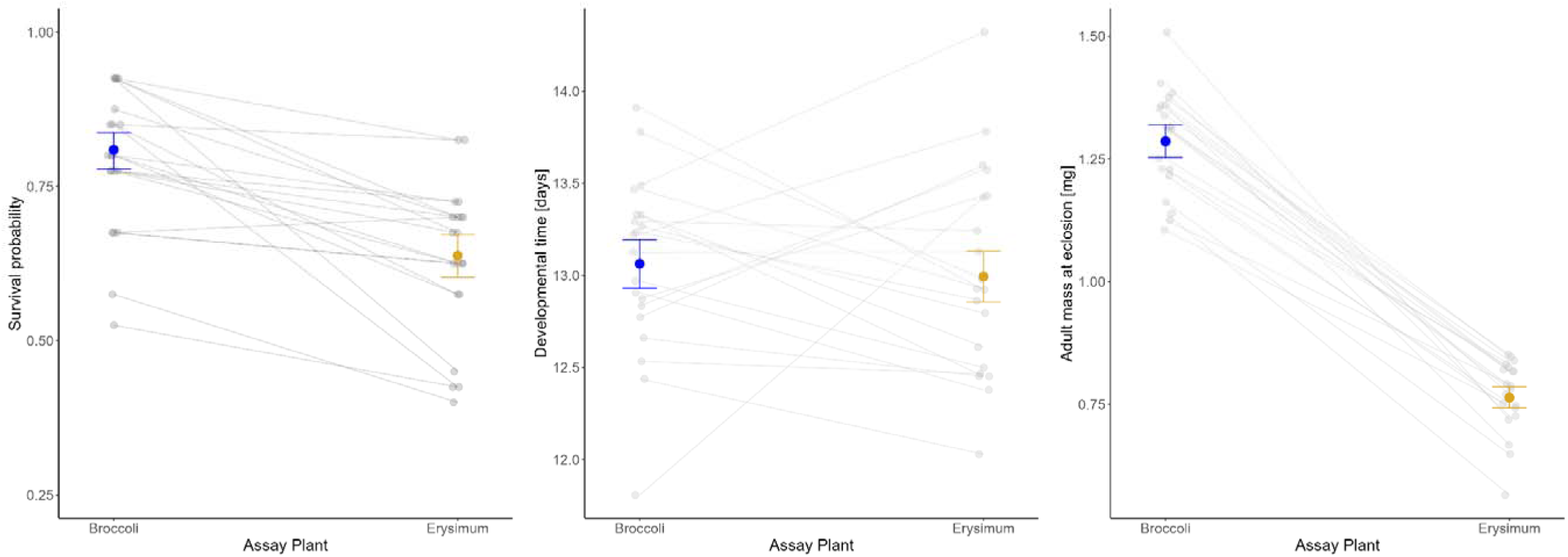
Life-history traits measured from the same life-history assays of 20 wild DBM populations: a) survival probability of second-instar larvae to adulthood; b) development time of female moths from assay setup to eclosion; and c) mass of female moths at eclosion. Results for male moths were fundamentally similar but are omitted for clarity. Coloured dots and error bars are global model-predicted marginal means with 95% confidence intervals, averaged across populations. Grey dots are means for each population, with grey lines connecting means of the same population on each assay plant.

Development time did not differ between assay plants (Figure 3b; χ² = 0.28, df = 1, p = 0.593), but differed strongly among populations (χ² = 63.56, df = 19, p < 0.001), with a significant interaction between population and assay plant (χ² = 62.65, df = 19, p < 0.001), and higher heritabilites for population means on each host plant (broccoli: H^2^ = 0.17; *Erysimum*: H^2^ = 0.26). Therefore, even though population means in development time had relatively similar ranges from 11.8 to 13.9 days on broccoli and 12.0 to 14.3 days on *Erysimum*, fastest-developing populations on broccoli were often among the slowest-developing populations on *Erysimum*, and vice versa (Figure 3b). We also found a main effect of sex on development time (χ² = 14.32, df = 1, p < 0.001), with males developing slightly slower than females (13.2 ± 0.05 *vs.* 13.0 ± 0.05 days).

Finally, adult mass at eclosion differed substantially with assay plant (Figure 3c, χ² = 50.62, df = 1, p < 0.001), with female moths on *Erysimum* reaching 40% lower mass compared to females on broccoli (0.76 ± 0.011 *vs.* 1.29 ± 0.017 mg). In addition, populations varied strongly in adult mass (χ² = 68.74, df = 19, p < 0.001), with population means ranging from 1.11–1.51 mg on broccoli and 0.56–0.85 mg on *Erysimum*. We found a significant interaction between population and assay plant (χ² = 57.80, df = 19, p < 0.001), with different populations growing fastest on broccoli and *Erysimum*, respectively (Fig. 3c), and modest heritability for population means on each host plant (broccoli: H^2^ = 0.12; *Erysimum*: H^2^ = 0.13). We also found significant effects of sex on adult mass (χ² = 245.33, df = 1, p < 0.001), as well as a significant interaction between sex and assay plant (χ² = 7.72, df = 1, p = 0.005). Male moths weighed 23% less than females on broccoli, whereas sexual dimorphism was reduced on *Erysimum*, with males weighing only 18% less.

Even though wild DBM populations differed significantly for all assays, we only found few significant correlations among population means for different choice and performance traits (Figure S2). For life-history assays, which yielded several traits, results substantially differed for within-plant and between-plant comparisons. Within assay plant, population means of development speed (inverse of development time) was positively correlated with adult mass at eclosion (broccoli: *r* = 0.65, p = 0.002; *Erysimum*: *r* = 0.69, p < 0.001), whereas between-plant, development speed and adult mass on broccoli vs. *Erysimum* were uncorrelated (Figure S2). Thus, while some populations always achieved fast development and high adult mass, the identity of best-performing populations changed between assay plants. In contrast, mean survival on the two assay plants was positively correlated with each other, but mostly not correlated with other life history traits. Instead, survival was negatively correlated with larval choice (Figure S2).

### 3.2. Continuous change in adult DBM mass during experimental evolution

We exposed replicate DBM lines to experimental evolution and monitored the mass of male and female adult moths in each generation (Figure 4). We found a strong effect of generation as a factorial variable (χ² = 1554.15, df = 23, p < 0.001), as well as an interaction between generation and selection regime (χ² = 337.11, df = 23, p < 0.001), suggesting a significant, treatment-specific change in mass over time. Even though there was substantial among-generation variation in adult mass, as is common in experimental evolution studies, the overall temporal trend was approximately linear (Figure 4), thus we simplified the model by specifying generation as a continuous variable, which did not affect overall results (generation: χ² = 1104.78, df = 1, p < 0.001; generation × selection regime: χ² = 107.24, df = 1, p < 0.001). Both models also supported an effect of sex (χ² = 358.54, df = 1, p < 0.001) and an interaction between sex and selection regime (χ² = 45.29, df = 1, p < 0.001).

**Figure 4.**
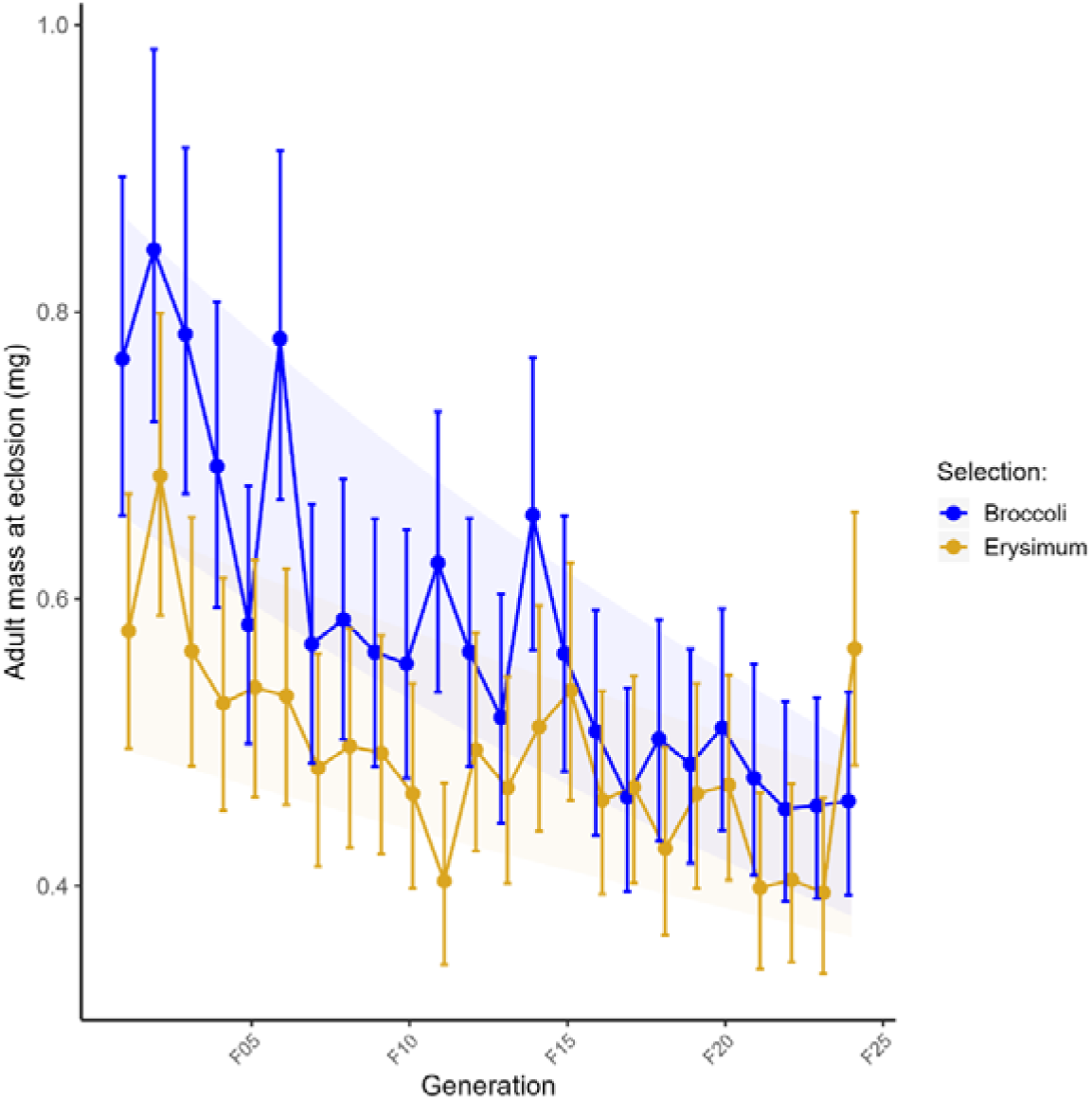
Mean mass of newly eclosed female moths, developing in broccoli (blue) or *Erysimum* (orange) over 24 generations of experimental evolution. Results for male moths were fundamentally similar but are omitted for clarity. Coloured dots are model-predicted marginal means with 95% confidence intervals from a model of log-transformed weights, treating generation as factorial variable. In addition, shaded areas are the 95% confidence interval from an equivalent model treating generation as continuous variable.

From the linear change model, it became obvious that adult mass differed substantially between selection regimes in the first generation (pairwise linear contrasts for females: t-ratio = 6.45, df = 1267, p < 0.001; males: t-ratio = 4.25, df = 1268, p < 0.001), with females and males initially weighing 25% and 17% less on *Erysimum* relative to broccoli, respectively. However, over the course of selection, adult mass in both treatments decreased significantly (log-slopes [95% confidence intervals]; broccoli: -0.0239 [-0.0253, - 0.0225]; *Erysimum*: -0.0133 [-0.0147, -0.0118]), with a significantly stronger decrease in mass for broccoli selection lines compared to *Erysimum* selection lines (pairwise linear contrast on slopes: t-ratio = -10.36, df = 5198, p < 0.001). As a result of this change, by generation 25 adult mass no longer differed between selection regimes (Figure 4; pairwise linear contrasts for females: t-ratio = 0.68, df = 1327, p = 0.495; males: t-ratio = -1.61, df = 1328, p = 0.108). Importantly, adult mass was unrelated to population size (Figure S3; correlation between mean female mass and mean adult number: *r* = 0.046, p = 0.757), thus this effect was unlikely caused by changes in crowding. Population size varied strongly over time (generation × selection regime: χ² = 65.66, df = 24, p < 0.001), but while population sizes were higher initially and underwent a reduction in generations 4 to 8, they stabilized to approximately 1000 individuals from generation 9 until the end of the experiment (Figure S3).

### 3.3. Change in preference and performance of DBM populations in response to selection

We performed standardized assays at different times during experimental evolution. Comparing larval preference in generations 3 and 26, we found that first-instar larvae from selection lines consistently avoided *Erysimum* (Figure 5). While selection regime (χ² = 0.20, df = 1, p = 0.657) and generation (χ² = 0.88.23, df = 1, p = 0.347) had no significant effect on preference, we found a marginally significant interaction between generation and selection regime (χ² = 2.96, df = 1, p = 0.085), with larvae from *Erysimum* selection lines showing slightly stronger avoidance of *Erysimum* compared to broccoli selection lines (Figure 5; pairwise linear contrasts: z-ratio = 1.94, p = 0.052).

**Figure 5.**
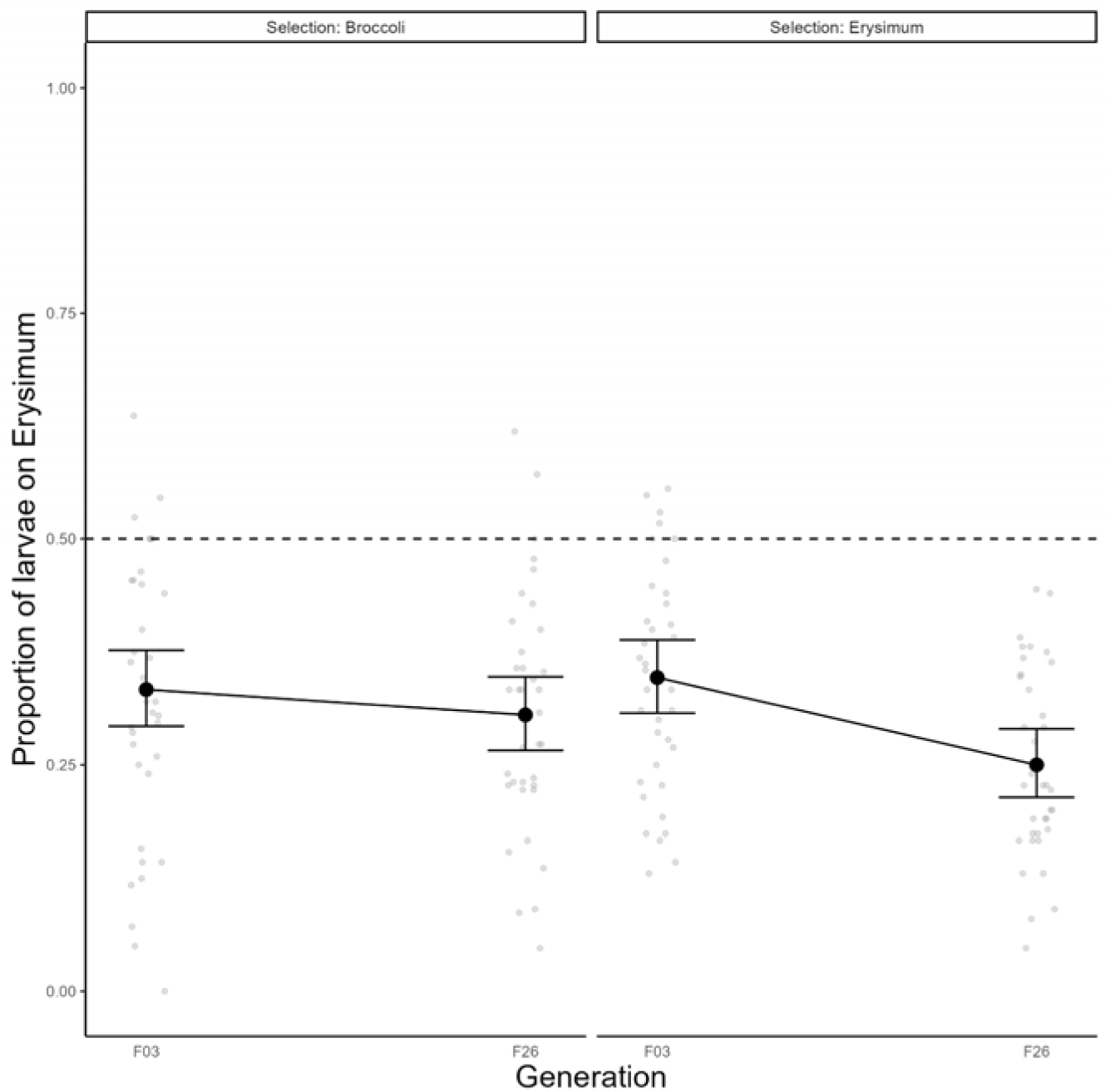
Choice of first-instar DMB larvae from selection lines feeding on *Erysimum* vs. broccoli leaves in two-way choice assays at generations 3 and 26 of experimental evolution. The left half of the figure shows selection lines evolving on broccoli (‘controls’), while the right half shows selection lines evolving on *Erysimum*. Black dots and error bars are the global predicted means with 95% confidence intervals, while small grey dots are replicate choice assays of approximately 24 larvae each.

Growth rate assays were performed in generations 3, 10, and 26, and similar as for wild DBM populations, we found strong effects of treatment in both selection lines across generations (Figure 6; χ² = 12.77, df = 2, p =0.002). Additionally, we found a marginal effect of selection regime (χ² = 3.78, df = 1, p = 0.052), as well as an effect of generation (χ² = 92.46, df = 2, p < 0.001) and an interaction between generation and treatment (χ² = 29.73, df = 4, p < 0.001). In generation 3, we found that the addition of digitoxin to broccoli reduced RGR_max_ by 2%, while feeding on *Erysimum* leaf discs reduced growth by 9%. By generation 10, these effects increased for both treatments (broccoli + digitoxin: -11%; *Erysimum*: -24%), while by generation 26, the differences increased disproportionately for larvae feeding on *Erysimum* leaf discs (Figure 6; broccoli + digitoxin: -11%; *Erysimum*: -37%). Overall, larvae from *Erysimum* selection lines had higher growth rates compared to broccoli lines (broccoli: 0.030 ± 0.001 mg mg^-1^ h^-1^; *Erysimum*: 0.033 ± 0.001 mg mg^-1^ h^-1^), but we found no support for significant change in this response as a result of experimental evolution (treatment × selection regime × generation: χ² = 2.85, df = 4, p = 0.582).

**Figure 6.**
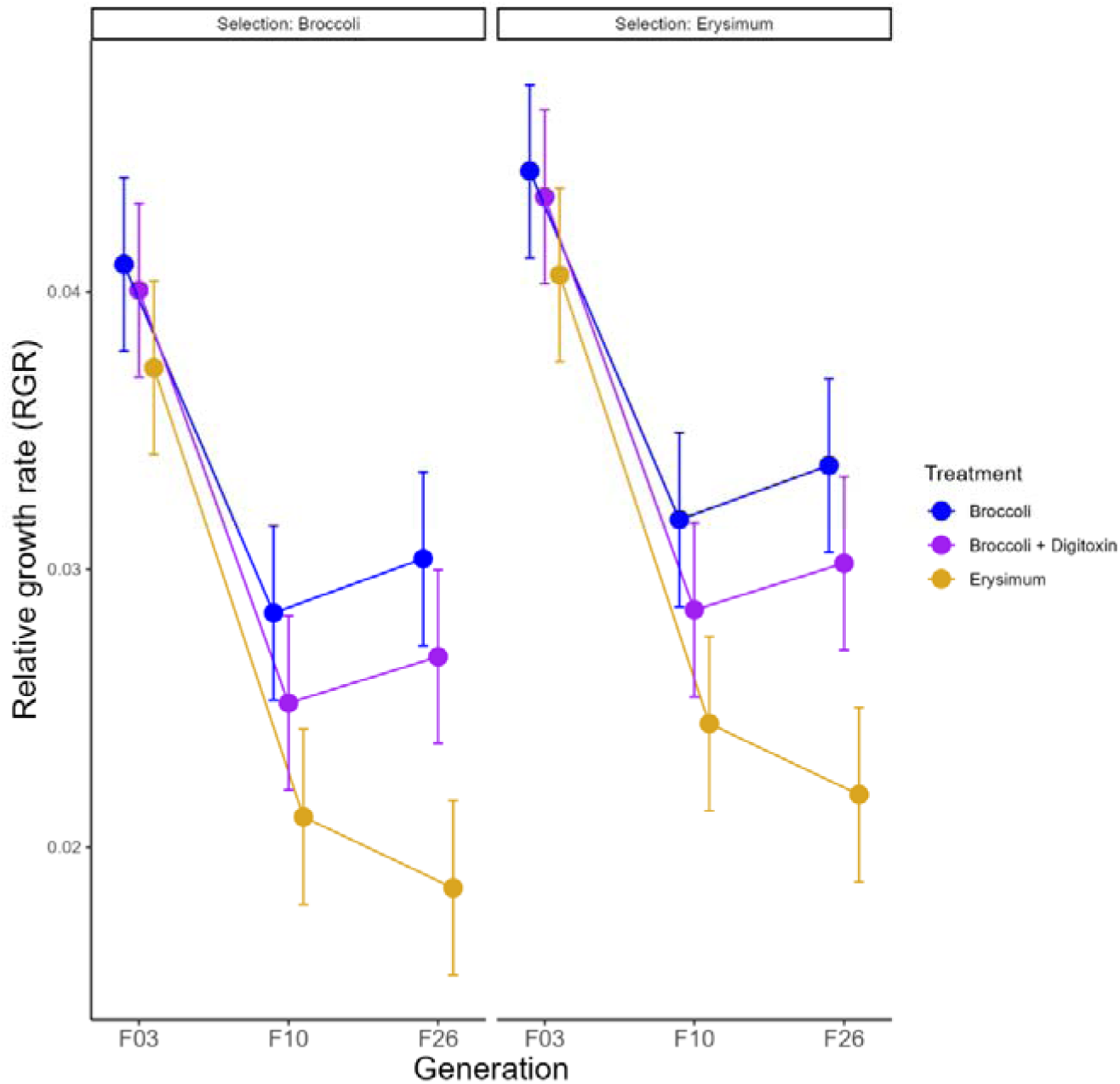
Growth of larvae from DBM selection lines at generations 3, 10, and 26 of experimental evolution. The left half of the figure shows selection lines evolving on broccoli (‘controls’), while the right half shows selection lines evolving on *Erysimum*. Coloured dots and error bars are global means of maximum RGR with 95% confidence intervals for larvae feeding on leaf discs of broccoli (blue), broccoli with added digitoxin (purple), or *Erysimum* (orange).

Life-history assays were performed most frequently, in generations 4, 12, 18, and 25. Survival from second-instar larvae to adulthood was strongly affected by assay plant (χ² = 50.29, df = 1, p < 0.001), with substantially lower survival on *Erysimum* (Figure 7a; broccoli: 0.715 ± 0.021; *Erysimum*: 0.511 ± 0.023). Survival additionally varied substantially between generations (χ² = 46.83, df = 3, p < 0.001), and while selection regime had no significant effect (χ² = 2.16, df = 1, p = 0.142), we found a significant interaction between generation and selection regime (χ² = 12.18, df = 3, p = 0.006).

**Figure 7.**
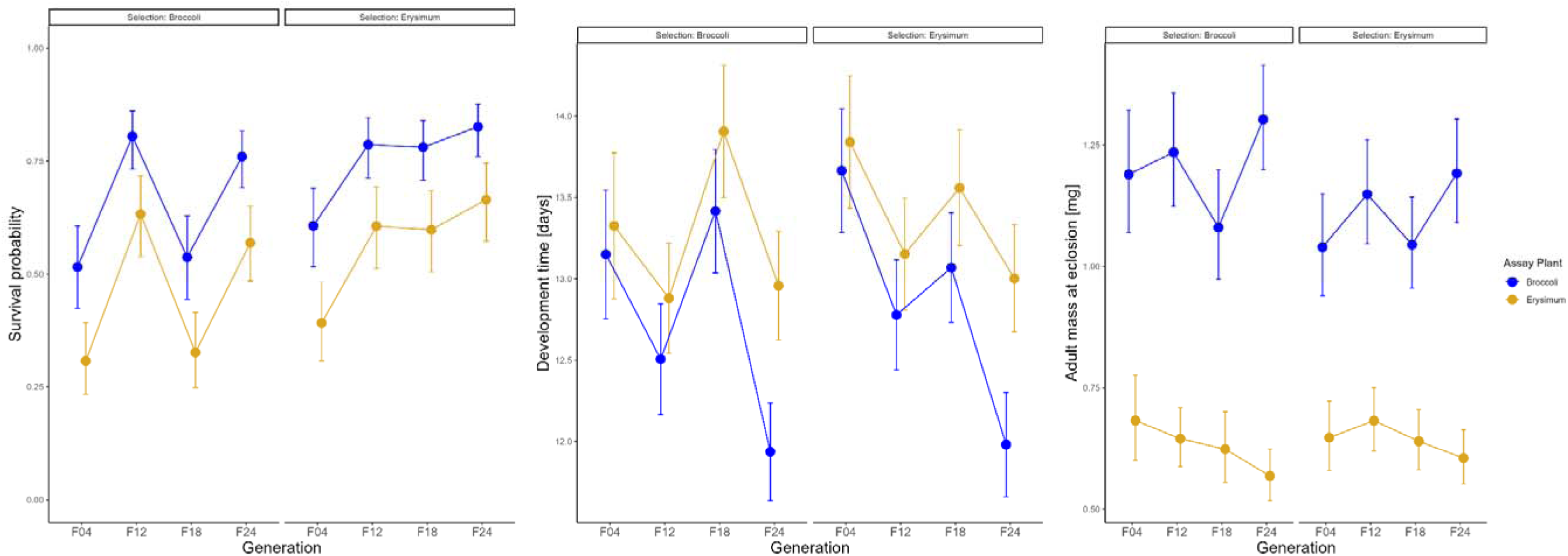
Life-history traits measured from the same life-history assays of DBM selection lines at generations 4, 12, 18, and 24 of experimental evolution: a) survival of second-instar larvae to adulthood; b) development time of female moths from assay setup to eclosion; and c) mass of female moths at eclosion. Results for male moths were fundamentally similar but are omitted for clarity. The left half in each panel shows selection lines evolving on broccoli (‘controls’), while the right half shows selection lines evolving on *Erysimum*. Coloured dots and error bars are global model-predicted marginal means with 95% confidence intervals, averaged across populations. Grey dots are means for each population, with grey lines connecting means of the same population on each assay plant.

Development time of larvae varied with selection regime (Figure 7b; χ² = 4.69 df = 1, p = 0.030) and generation (χ² = 46.35 df = 3, p < 0.001). Additionally, we found support for interactions of generation with assay plant (χ² = 9.89.18, df = 3, p = 0.020) and with selection regime (χ² = 8.32, df = 3, p = 0.040). Overall, all larvae accelerated their development in response to selection, and while generation 18 deviated from this pattern, at the end of the selection experiment, larvae from either selection regime completed development over one day faster when feeding on broccoli. Interestingly, while females from *Erysimum* selection lines initially developed slower than broccoli lines on either assay plant (generation 4, pairwise linear contrasts: t-ratio = -2.15, p = 0.033), these differences disappeared by the end of selection experiment (generation 24: t-ratio = -0.24, p = 0.815).

Finally, adult mass at eclosion was affected by assay plant (Figure 7c; χ² = 94.74, df = 1, p < 0.001), with female moths on *Erysimum* reaching on average 45% lower mass compared to females on broccoli (0.635 ± 0.013 *vs.* 1.51 ± 0.022 mg). Additionally, we found support for interactions of assay plant with generation (χ² = 18.89.15, df = 3, p < 0.001) and with selection regime (χ² = 5.90, df = 1, p = 0.015). Interestingly, adult mass of individuals developing on *Erysimum* assay plants slightly decreased regardless of selection regime (generation 4 vs. 24, pairwise linear contrast: t-ratio = 2.30, p = 0.022), whereas adult mass increased for individuals feeding on broccoli plants (Figure 7c; generation 4 vs. 24, pairwise linear contrast: t-ratio = -2.58, p = 0.011).

## 4. Discussion

We intensively sampled naturally occurring DBM populations in Switzerland and uncovered substantial variation in behavioural and performance traits for larvae developing on cardenolide-containing *Erysimum* plants. We show that young larvae avoided feeding on the *Erysimum* leaves, while older larvae suffered reduced growth, had reduced survival, and achieved substantially lower adult mass. Nonetheless, wild populations feeding on *Erysimum* showed substantial trait variation, with heritability estimates indicating that genetic differences accounted for a meaningful proportion of variation in *Erysimum* avoidance, development time, and adult mass. Using experimental evolution, we therefore attempted to select for and disentangle these components, but even after 25 generations of selection, *Erysimum*-evolved lines remained indistinct from broccoli-evolved control lines in their performance on *Erysimum*. Instead, all selection lines converged on one phenotype regardless of selection regime, providing important insights into the selective landscape faced by herbivores on a novel host plant.

First-instar larvae of wild DBM populations had an innate avoidance of *Erysimum* leaves, and this avoidance behaviour remained after 25 generations of selection. In fact, *Erysimum*-evolved selection lines even became marginally more repelled by *Erysimum* leaves compared to broccoli-evolved lines. Cardenolides have been shown to be effective feeding deterrents for several other insect species (Akhtar & Isman, 2003; Green et al., 2011; Petschenka et al., 2011). For *Erysimum*, Dimock et al. (1991) demonstrated that larvae of *Pieris rapae* refused to feed on leaves, with cardenolides being identified as the main repellent compounds (Dimock et al., 1991; Sachdev-Gupta et al., 1993). Similarly, cardenolides were shown to prevent oviposition on *Erysimum* plants by adult *Pieris rapae* butterflies (Renwick & Radke, 1987; Rothschild et al., 1988), a finding which was recently confirmed using *Erysimum* CRISPR-Cas9 plants with abolished cardenolide production (Younkin et al., 2024). The perception of cardenolides likely involves taste receptors and may use a balance of feeding stimulants and deterrents in decision making (Kasubuchi et al., 2018). Our results cannot confirm a causal role of cardenolides for the observed avoidance of *Erysimum* by DBM, and at least for older, third instar DBM larvae, cardenolides appear to have limited effects on choice behaviour (Wang & Züst, 2025). Nonetheless, avoidance of *Erysimum* had the highest heritability of all our measured traits, with mean avoidance by DBM populations remaining consistent over several generations, thus a potential involvement of taste perception of cardenolides by first-instar DMB larvae should be confirmed in future experiments.

Late second-instar larvae of wild DBM populations feeding on *Erysimum* leaves gained less weight compared to larvae feeding on control broccoli plants, resulting in a reduction in growth rates of approximately 20%. In these controlled growth rate assays, the addition of digitoxin impacted DBM weight gains, but only resulted in a growth rate reduction of approximately 10%. This suggests that cardenolides may play a partial role in reducing DBM growth on *Erysimum*, but it also indicates that other *Erysimum* traits are likely to be important as well. Cardenolides have been shown to impair growth of larvae in other lepidopteran systems; for example, larvae of *Trichoplusia ni* exhibited increased growth on cardenolide knockout plants (Younkin et al., 2024), and cardenolides added to diet or directly administered orally reduced growth of *T. ni*, *Lymantria dispar*, and *Bombyx mori* (Dussourd & Hoyle, 2000; Fukuyama et al., 1993; Karowe & Golston, 2006). Nonetheless, these effects likely are not universal for all insects, while other traits, such as the dense trichomes of *Erysimum*, may be more important for the defence against DBM (Wang & Züst, 2025).

Larval growth rates decreased substantially over successive generations under both selection regimes and for all assay treatments, with the strongest decrease observed on *Erysimum* leaves. It is important to note that this decrease is exaggerated by very high mean larval growth rates in generation F03, which were higher even than most population means of wild DBM ancestors. It is possible that this was caused by a certain amount of ‘hybrid vigour’ which then dissipated over subsequent generations, an effect which has been demonstrated for termite colonies (Lee et al., 2020). However, we did not observe equivalent effects for other performance traits, while larval growth rates also had low heritability among wild DBM ancestors. This suggests high environmental plasticity or stochasticity in this trait, which should therefore be interpreted with caution.

Larvae of wild DBM populations had lower survival and reached substantially lower adult mass when developing under controlled, low-density conditions on *Erysimum*, compared to larvae developing on broccoli. Mean survival exhibited no heritability among wild DBM ancestors and was therefore primarily the result of environmental or stochastic processes. Consequentially, survival rates were somewhat variable in response to selection but remained consistently lower on *Erysimum* relative to broccoli. In contrast, both development time and adult mass exhibited larger heritability and both traits showed consistent changes in response to selection, albeit not in a regime-specific fashion. Larvae from both selection regimes accelerated their development time on broccoli over the course of selection, increasing the differences between performance on assay plants over time. Similarly, adults in low-density broccoli assays generally grew larger in successive assays, while adult mass on *Erysimum* assay plants continuously decreased.

Interestingly, mean development time and adult mass on *Erysimum* after 25 generations of selection approximately matched the lowest values found among population means of wild DBM ancestors, suggesting that both selection regimes favoured a phenotype already present at the onset of selection, with fast development but low mass on *Erysimum*. In contrast, individuals achieving higher mass on *Erysimum* were apparently not retained by selection. Even though adult mass in insects is expected to directly relate to lifetime fecundity (Honěk, 1993) and should therefore be adaptive, it may also be related to other traits such as flight musculature (Shirai, 1995). It is thus feasible that under the specific conditions of our experiment that lacked a requirement for dispersal, higher mass may not have provided a strong fitness benefit.

In contrast to the relatively poor performance on *Erysimum*, larvae evolving on broccoli accelerated their development while also slightly increasing their mass at eclosion. Development time of insect larvae is often a plastic response that can rapidly evolve in response to directional selection, but accelerated development often comes at the cost of reduced adult mass due to physiological trade-offs (Mueller, 1988). In our study, we found no evidence for such trade-offs and instead found that development speed and adult mass were already positively correlated among our sampled wild DBM populations. Perhaps even more surprisingly, the mean traits of evolved populations from both selection regimes closely matched the traits of individual ancestral populations. For example, larvae from population FR.KERZ developed at the same speed and reached even higher adult mass as evolved lines on broccoli, while they developed at slower speeds and reached lower mass comparable to evolved lines on *Erysimum*.

The relatively modest change in traits over selection under controlled, low-density assay conditions was in stark contrast to the changes in mass of adults directly emerging from populations under selection. Here, adults emerging from broccoli selection lines were initially much heavier than adults emerging from *Erysimum* selection lines. However, over successive generations of selection, adult mass decreased almost linearly, resulting in adults in the final generation weighing almost half as much as in the first, and for adults from the two selection regimes no longer differing in mass. Adult mass could be influenced by larval density and crowding, but the initial three generations experienced the highest densities and nonetheless achieved the highest mass, while densities stabilized for later generations, and adult mass nonetheless continued to decline. Even though we do not have estimates of effective population sizes, each selection line was maintained at a high number of individuals throughout. While numbers fluctuated somewhat, only few populations dropped below 200 individuals, which is unlikely to represent a genetic bottleneck, and results are therefore unlikely to be the result of drift alone. Therefore, these results are again more in line with selection for a highly plastic DBM phenotype by both selection regimes. The favoured phenotype grows rapidly to small adult sizes under suboptimal, crowded conditions, but retains the ability to grow larger under optimal, low-density conditions. This would suggest that developing on *Erysimum* even at low densities poses similar limitations on DBM growth as developing on crowded, heavily attacked broccoli plants.

The nature of selection imposed on DBM by developing on *Erysimum* is a key point in understanding its evolutionary responses and may explain why *Erysimum*-evolved DBM lines failed to improve their performance on *Erysimum*. Our results are comparable to those of Zalucki et al. (2021), who maintained the generalist lepidopteran herbivore *Helicoverpa armigera* on a glucosinolate-producing host and after 30 generations of selection found no evidence of improved performance. Despite their fundamental differences in modes of activity, coping with glucosinolates in the absence of specialized adaptations likely involves similar mechanisms as coping with cardenolides, i.e., modification and excretion (Jeckel et al., 2022). Clearly, both DBM and *H. armigera* can complete development on their respective novel hosts but apparently lack the type of standing genetic variation required for substantially improving performance. At least for DBM on *Erysimum*, overcoming cardenolides may only be a first barrier, while other barriers, such as coping with physical defences (trichomes) or low nutritional quality may be harder to overcome. Therefore, even though experimental evolution can result in rapid change from standing variation (e.g., Ramos & Schiestl, 2019; Züst et al., 2012), they have clear limitations when novel traits are required to evolve. New methods will therefore have the potential to revolutionize the field, where ‘knock-in’ of resistance traits by CRISPR-Cas9 (Karageorgi et al., 2019) could be followed by experimental evolution.

Somewhat surprisingly, the prominent and potent cardenolide defences gained by *Erysimum* in evolutionary recent times do not appear to be strongly involved in DBM resistance. How DBM copes with cardenolides, and which other traits are primarily responsible for its low performance on *Erysimum* remains an open question. Nonetheless, the current baseline tolerance of *Erysimum* by DBM could well represent an evolutionary stepping-stone for increased tolerance should *Erysimum* become more abundant in the future, but the required traits may be entirely unrelated to cardenolide tolerance.

Overall, our study provides insights into the complex ecological and evolutionary dynamics of the interaction between unadapted herbivores and novel host plants. Overcoming of potent toxic defences is often seen as a key limitation for herbivores to use new hosts, but our results reveal that chemical traits may commonly be combined in suites of other defence traits that may even outweigh chemical traits in their importance in preventing herbivore attack. Our view of the co-evolutionary arms race that matches a chemical plant defence with an herbivore offence strategy thus is almost certainly an idealized state, biased by a predominantly retrospective analysis of co-evolved systems. Prospective analysis of systems currently undergoing coevolution, such as our study, therefore promises a more unbiased view of the likely outcome of co-evolutionary interactions.

## Acknowledgements

We thank Rayko Jonas and Markus Meierhofer for invaluable support in the production of feeding plants, and Chiara Lardi and Lucia Salis for their assistance with maintenance of DMB populations. Many farmers kindly granted us access to their fields during collection of wild DMB. This project has received funding from the European Research Council (ERC) under the European Union’s Horizon 2020 research and innovation programme (grant agreement No 950319). Additional support was provided by a Swiss National Science Foundation grant PCEFP3_194590 to TZ and a grant of the Georges and Antoine Claraz Foundation.

## Supplementary Materials

**Figure S1.**
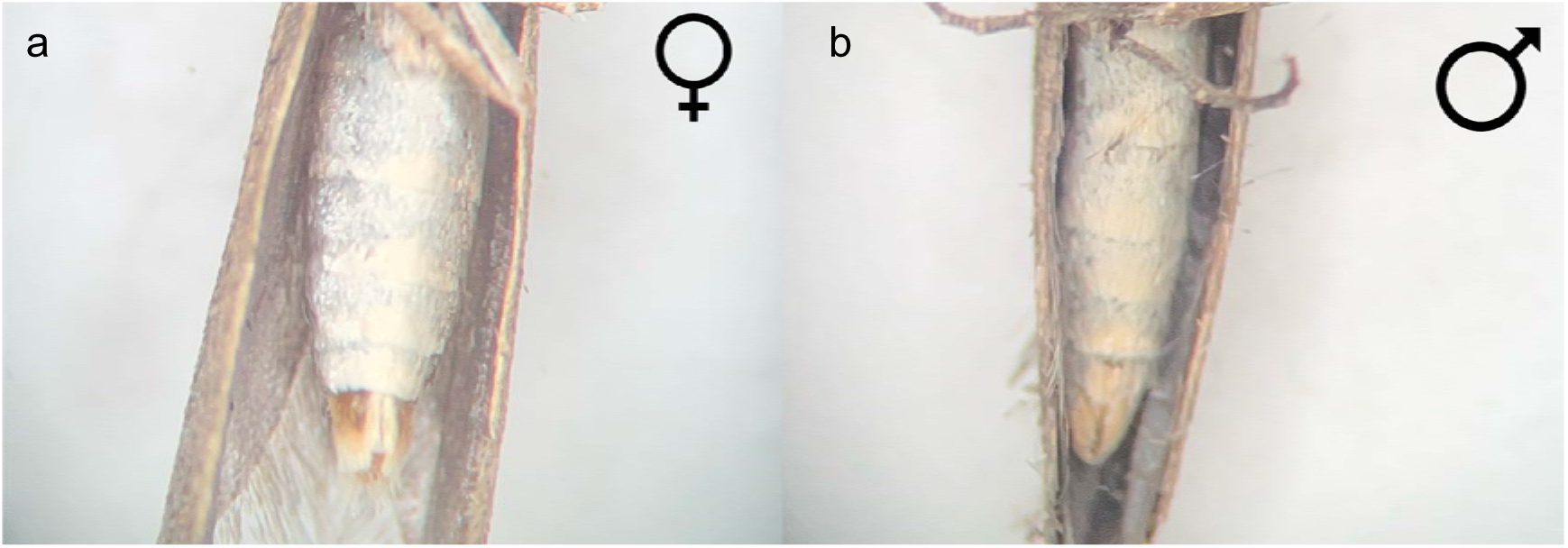
Female (a) and male (b) abdomen of adult *Plutella xylostella* moths. The shape of the final segment was used for sex determination.

**Figure S2.**
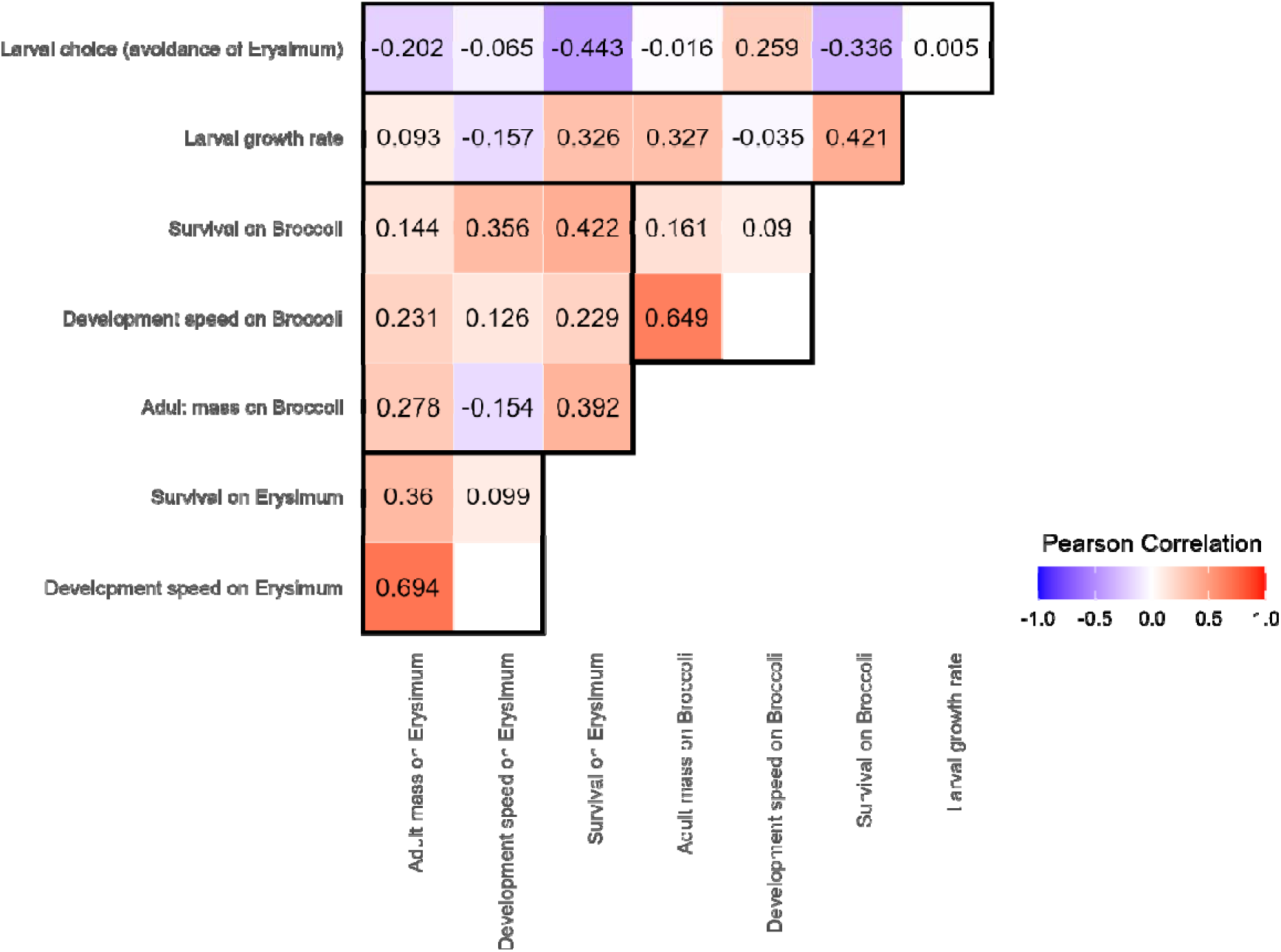
Correlations of mean preference and performance traits of 20 wild DBM populations measured in association assays, growth-rate assays, and life-history assays. Larval choice is summarized as ‘avoidance of *Erysimum’* (inverse proportion of larvae found on *Erysimum*), with higher values corresponding to stronger avoidance. For larval growth rate, no interaction between population and treatment was found, thus the relative ranking of populations remained the same regardless of host plant. Life-history assays produced three traits that varied substantially between host plant. For each trait, higher values correspond to better performance (‘development speed’ is the inverse of development time), while lower or more negative values correspond to worse performance. Heatmap colours correspond to correlation strength. Black squares or rectangles group together assays, with life-history traits are further grouped as within- and across-plant comparisons.

**Figure S3.**
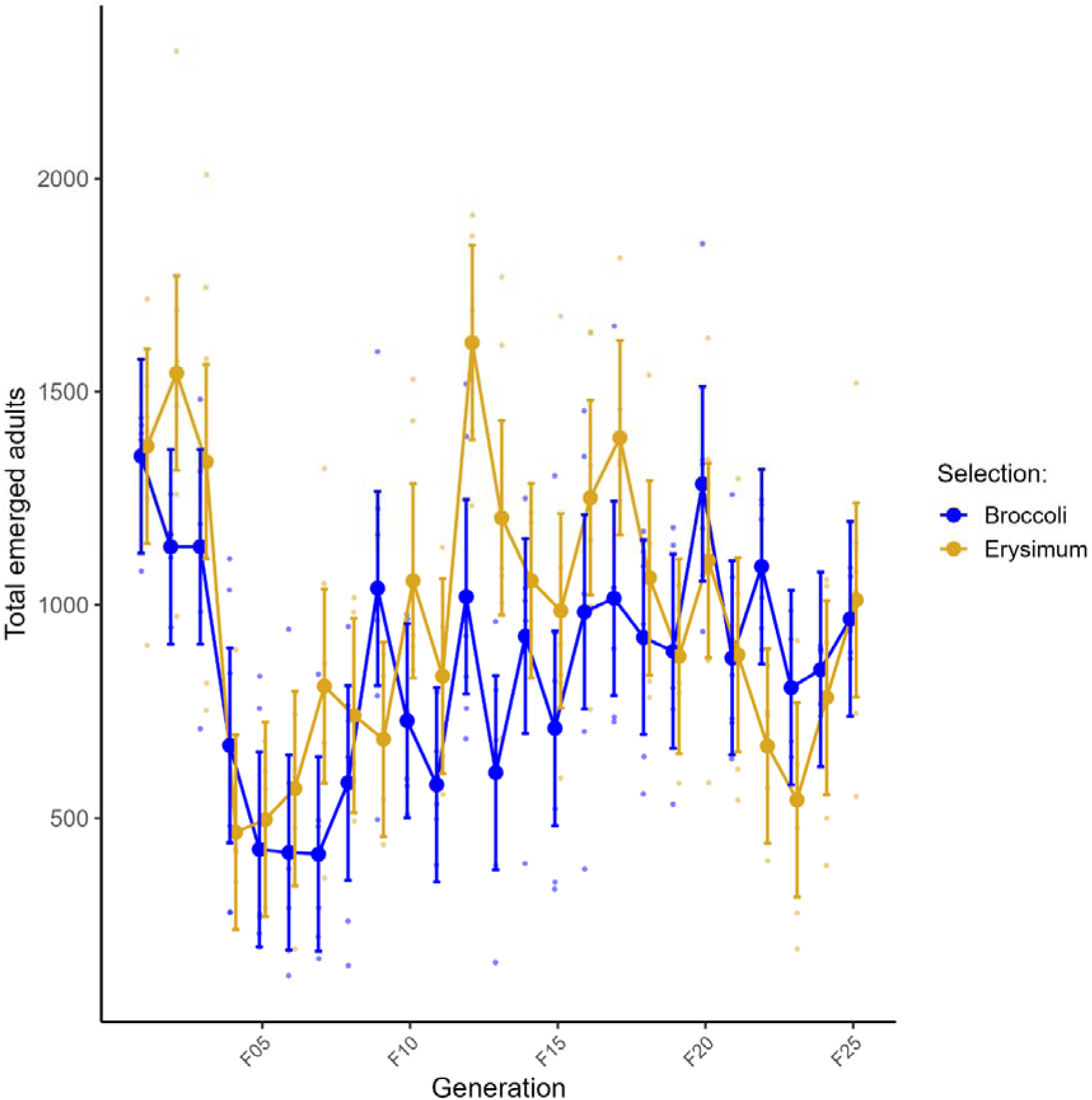
Mean total number of adult DBM emerging from selection lines on broccoli (blue) or *Erysimum* (orange) over 24 generations of experimental evolution. Large coloured dots and error bars are model-predicted marginal means with 95% confidence intervals from a model treating generation as factorial variable, while small dots are means for each replicate selection line.

**Table S1.**
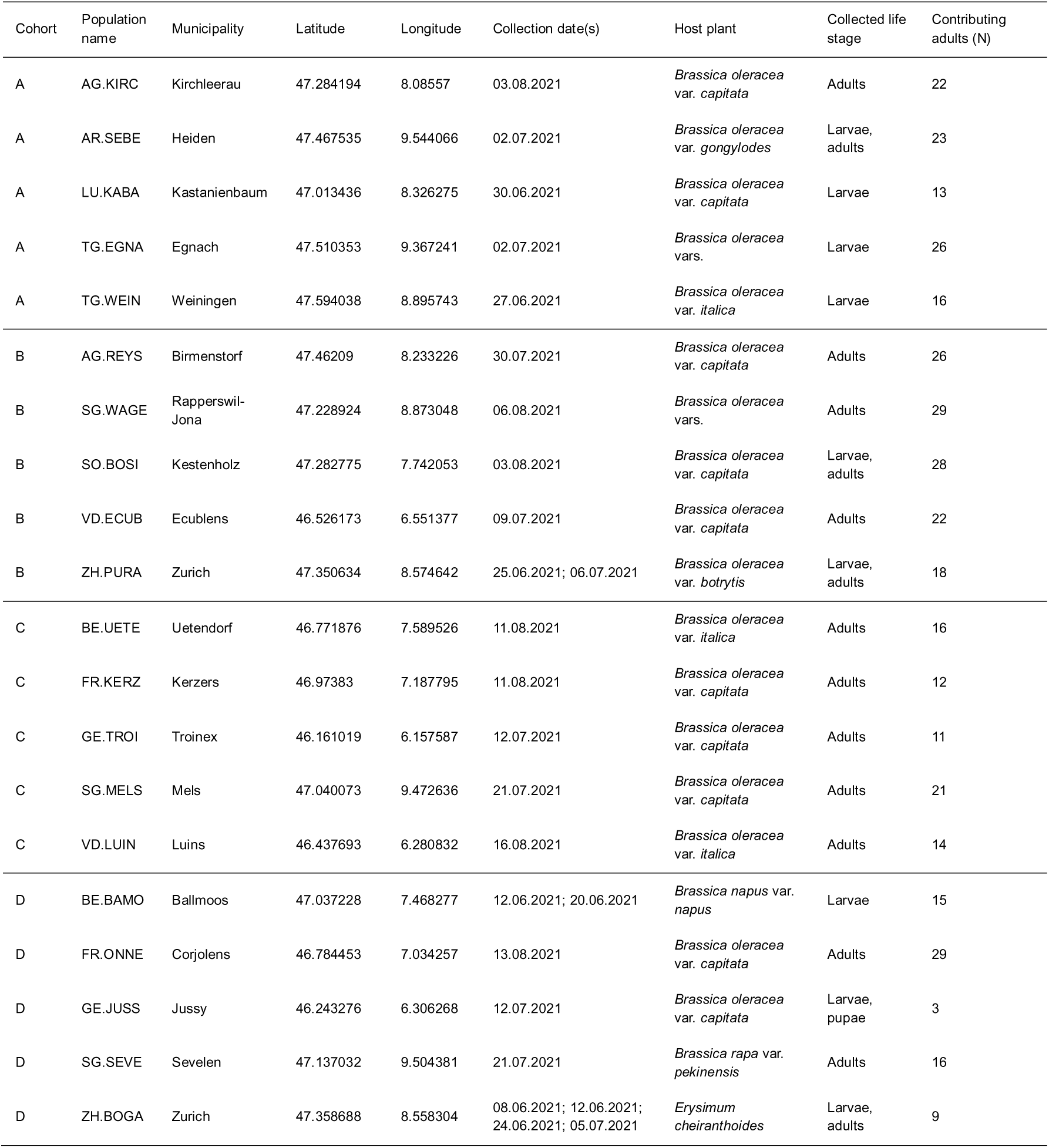
Collection locations for 20 wild populations of *P. xylostella*. Populations are grouped by rearing cohort and sorted alphabetically. Individuals were collected on one or several dates in 2021 from a variety of host plants growing in agricultural fields (except for ZH.BOGA). In each location the predominant life stage was collected in the field, and for larvae and pupae, reared to the adult stage in the laboratory. Due to mortality during transport and high parasitism rate of subadult stages, the initial population size N is approximated as the number of adults with the opportunity to contribute genes to the first generation.

**Table S2.**
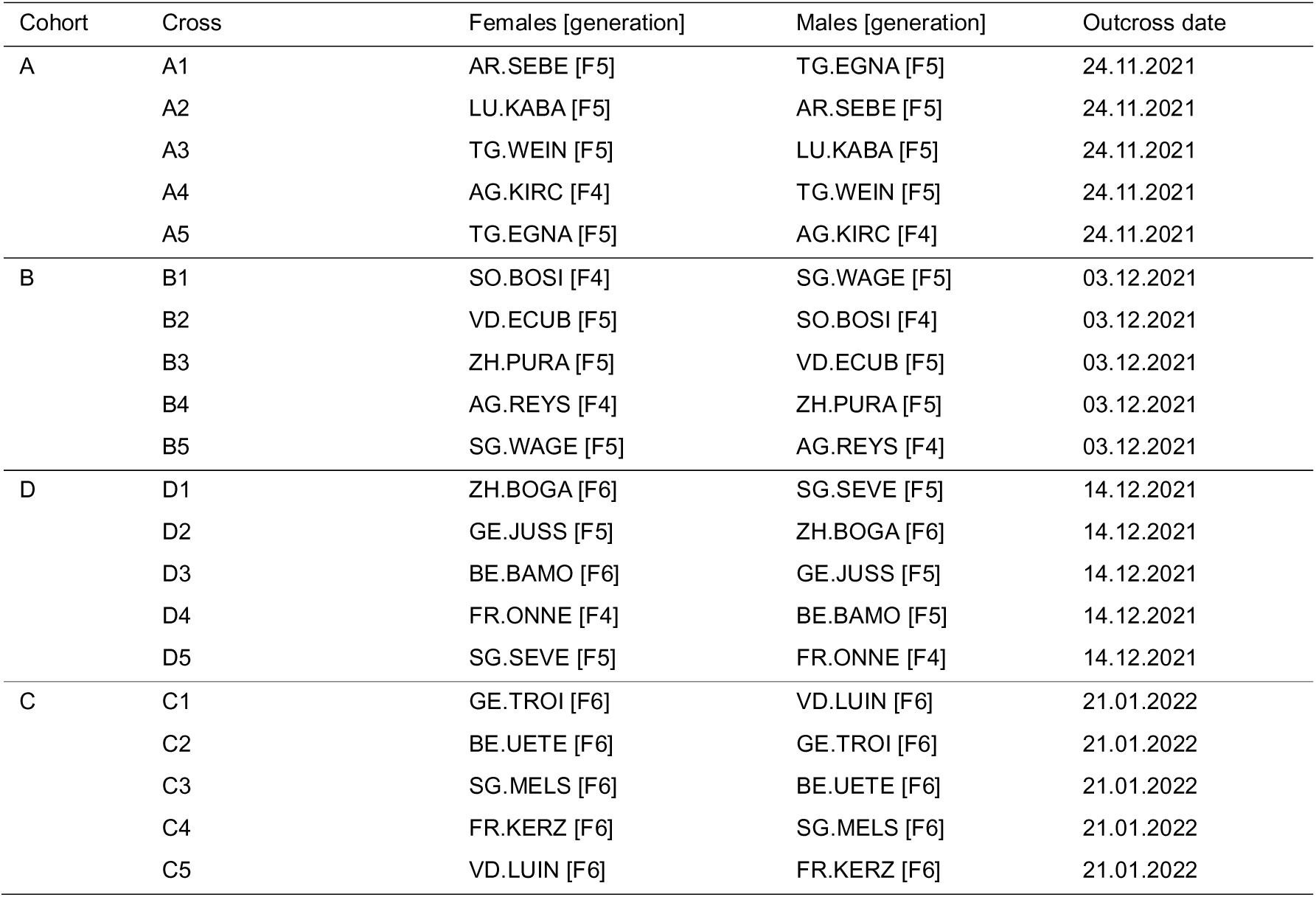
Crossing scheme for the first step in creating a panmictic population to be used for experimental evolution. Within each cohort, 30 females were combined with 30 males from another population, ensuring that each population was outcrossed with two other populations. At time of outcrossing, wild populations had been maintained in the laboratory for 4-6 generations, depending on time of collection.

